# RheoScale 2.0: Revealing the Hidden Roles of Protein Positions via Substitution Patterns

**DOI:** 10.64898/2026.08.10.743964

**Authors:** David H. Liu, Shwetha Sreenivasan, Carter J. Gray, Hannah C. Cleveland, Liskin Swint-Kruse

## Abstract

A central challenge in molecular biology is understanding how amino acid substitutions modulate various features of protein function and stability. To illuminate the complexities of this relationship, high-throughput (HTP) assays are increasingly used to assess site-saturating mutagenesis libraries. A common downstream analysis is to average the set of twenty outcomes at each amino acid position for comparison with structural and evolutionary features. Average values clearly identify positions that tolerate most substitutions (neutral positions) and positions where most substitutions abolish activity (toggle positions). However, average values conceal the existence of rheostat positions, where different amino acid substitutions sample a wide range of outcomes. To quantitatively identify rheostat positions, we previously developed a histogram-based analysis that we here expand by: (i) incorporating new position classes observed in experimental studies of rheostat positions; (ii) formalizing a hierarchy of class assignments; (iii) refining error-based identification of neutral positions; and (iv) statistically assessing the robustness of class assignments to changes in experimental and computational parameters. RheoScale 2.0 is implemented in Excel and newly implemented in Python for facile integration with existing HTP pipelines; all parameters are customizable. Example analyses are shown for three HTP datasets of the SARS-CoV-2 papain-like protease. Results illustrate two aspects that influence interpretation of HTP data: First, position assignments (and substitution outcomes) depend highly on the measured feature. Second, many protein positions play multiple roles in the sequence-structure-function relationship. The recognition of varied position roles will advance understanding of pathogen evolution, protein engineering, and variant interpretation for personalized medicine.

**Summary:** RheoScale 2.0 improves how high-throughput mutational data are interpreted by identifying protein positions where amino acid substitutions act like biological dimmer switches. By enabling more nuanced assignment of position behavior, beyond neutral or deleterious outcomes, this analysis framework advances studies of sequence-structure-function relationships and has broad relevance for understanding protein evolution, engineering proteins with desired properties, and interpreting variants linked to human disease.

**SOFTWARE AVAILABILITY:** https://github.com/liskinsk/RheoScale-calculator

## Introduction

Advances in personalized medicine and protein engineering, as well as recognizing which changes are significant for pathogen evolution, require understanding how variations in amino acid sequence impacts protein function and/or stability. Despite decades of effort, outcomes from single substitutions cannot yet be reliably predicted by computational algorithms (*e.g.,* ^1–5^). This challenge is, in large part, due to difficulties identifying biologically-meaningful substitutions at evolutionarily non-conserved positions and/or positions far from binding sites (*e.g.,* ^6–9^). Historically, these types of positions were overlooked in substitution experiments and, thus, under-represented in early computational training/test sets. To address this gap, over the past decade, numerous high-throughput (HTP) approaches have been developed to generate whole-protein measurements of substitution effects (*e.g.*, ^10–14^).

These HTP approaches yield a wealth of data but also raise analytical challenges: how can outcomes for multiple substitutions per position be meaningfully compared to structural features or evolutionary patterns? For these comparisons, a position-based analysis is required. One common approach uses each position’s set of substitution outcomes to calculate the average value. This metric effectively discriminates “toggle” positions, where most substitutions severely impair one or more protein features (*e.g.,* binding affinity, allosteric regulation), from “neutral” positions, where substitutions have minimal effects (Figure 1). However, average values might fail to discriminate “rheostat” positions (Figure 1, yellow histogram), where substitutions sample a wide range of outcomes. This oversight leads to missed opportunities for understanding the protein sequence-structure-function relationship.

**Figure 1.**
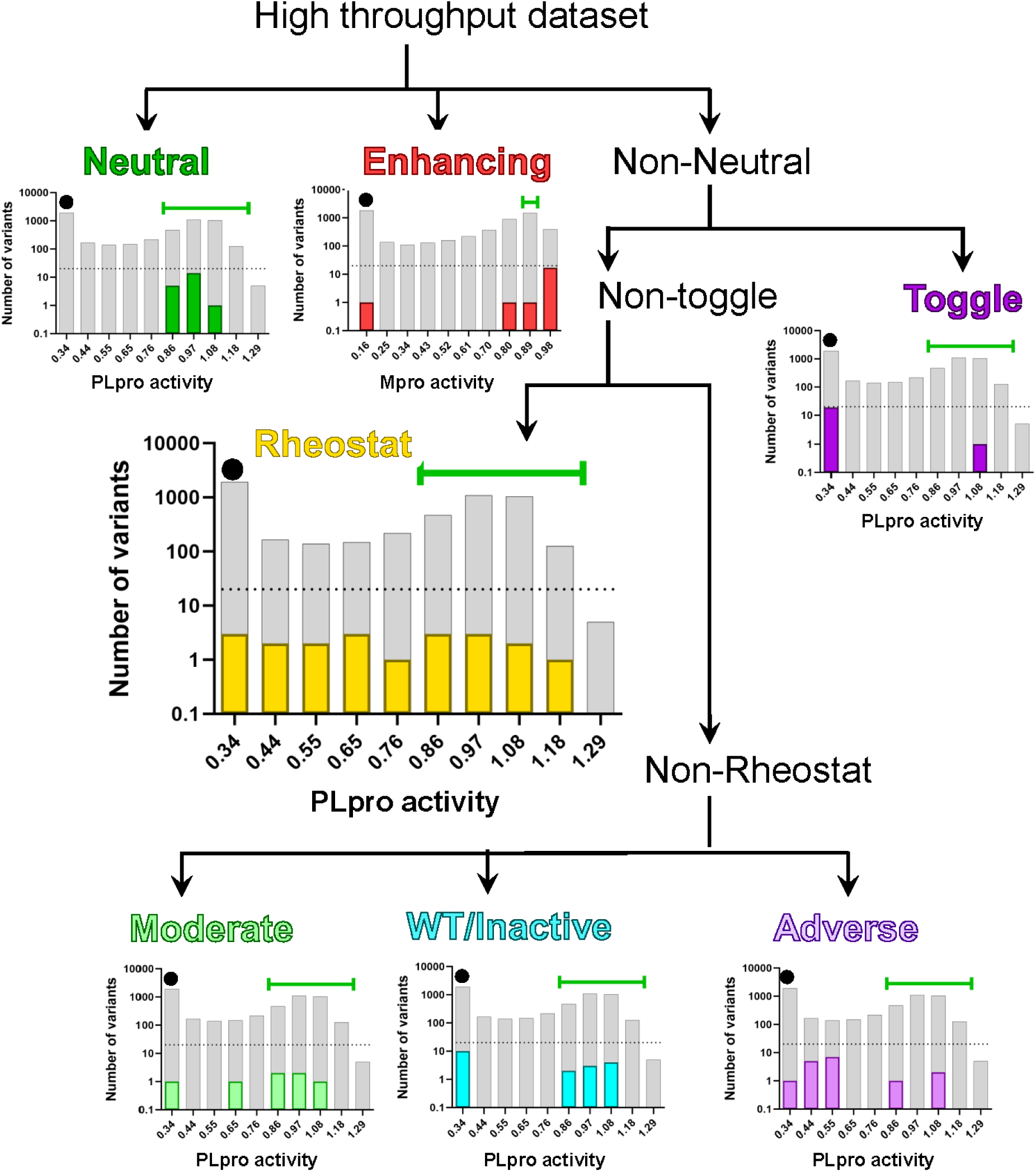
Hierarchical assignment of protein positions using RheoScale 2.0. HTP datasets are frequently presented as histograms. The gray histograms shown here represent the outcomes for all possible substitutions in SARS-CoV2 PLpro^12^ (or its Main protease, Mpro^10^). RheoScale 2.0 uses these histograms to assess the overall substitution sensitivities of individual positions. Analyses use three reference points: (i) the range of values observed for wild-type replicates (WT; green horizontal bars), (ii) the value corresponding to zero activity (black dot on the left-most bar), and (iii) a value for the “most active” end of the range, which is usually determined from empirical observations. Activity values on the x-axes correspond to the upper limits of each histogram bin, in the logarithmic units reported by the studies’ authors. The y-values are the number of variants observed and are also shown on log scale. Each position’s assignment is based on the pattern observed for its set of substitution outcomes. Assignments are made hierarchically as shown. Examples for each assignment are: Neutral (PLpro position 2); Enhancing (SARS-CoV-2 Mpro position 151); Toggle (PLpro position 109); Rheostat (PLpro position 82); Moderate (PLpro position 31), WT/Inactive (PLpro position 139), and Adverse (PLpro position 10). Parameters used to classify PLpro positions are provided in the example analysis files posted at github https://github.com/liskinsk/RheoScale-calculator; RheoScale analysis of Mpro was reported by Sreenivasan *et al.*^6^

Indeed, accumulating experimental evidence shows that rheostat positions are widespread.^7^ They occur in many types of proteins, including globular and intrinsically disordered transcription factors, globular enzymes, integral membrane transport proteins, and extracellular receptors. In several proteins, >30% of their positions exhibit rheostatic behavior.^6; 8; 15^ This observation is mirrored in analyses of proteins in the Swiss-Prot and EXPV datasets, where >30% of amino acid positions were predicted to be rheostat positions by the “funtrp” algorithm.^16^

Substitutions at rheostat positions can also modulate a wide variety of protein features, including: ligand binding, allosteric regulation, substrate transport, ligand specificity, and protein stability.^7^ Some rheostat positions simultaneously modulate multiple protein features.^7^ Substitution outcomes on function seldom correlate with amino acid physicochemical properties and show no consistent relationships to binding sites or solvent exposure.^7^ Instead, the structural regions around rheostat positions appear to be “plastic”, accommodating substitutions via localized changes.^7^ Other evidence suggests that substitutions at rheostat positions alter protein dynamics.^17–19^

Finally, many computational algorithms misclassify substitution outcomes at rheostat positions.^5^ This surely leads to missed-and mis-diagnoses in personalized medicine and impedes our understanding of pathogen evolution. A better understanding of rheostat positions would also advance protein engineering: (i) Rheostat positions provide ideal targets for fine-tuning various protein features, such as activity, ligand specificity, and/or stability and (ii) substitutions at rheostat positions often exhibit gain-of-function outcomes.^7^ For example, changes at rheostat positions were critical for optimizing the functions of novel chimeric transcription factors,^20; 21^ and a gain-of-function substitution at a rheostat position in a bile acid transport protein is a medically-relevant polymorphism.^22^

Clearly, it is vital to objectively identify rheostat positions in experimental datasets for further study. To that end, we previously developed a histogram-based analysis to quantitatively distinguish rheostat, neutral, and toggle positions.^23^ Since then, we have identified new position classes,^6; 8; 24^ formalized a hierarchy for assigning position types, and improved error-based identification of neutral positions^25^. Here, we unify these advances into RheoScale 2.0 and statistically assess effects of experimental and computational parameters on position assignments. These new features are integrated into the Excel version of the calculator and newly implemented as a Python package for facile integration into HTP pipelines.

## Methods

### Calculator overview

For a protein’s set of single amino acid substitutions, RheoScale 2.0 uses a modified histogram-analysis of assay values to classify the roles of individual protein positions (Figure 1). To that end, the histogram for each position is used to calculate four distinct scores – neutral, rheostat, toggle, and enhancing; mathematical definitions for these scores are included in the instruction sheets that accompany the calculators and in ^5; 6; 8; 23; 25^. Each set of scores is then used to classify each position using the hierarchy described below. For these assignments, default score thresholds are based on prior experimental studies; for bespoke analyses, override features are available for all parameters, as discussed further below.

RheoScale 2.0 is available on GitHub in two formats:

1. An interactive Excel workbook. A “small” version of the workbook analyzes up to 19 substitutions for up to 20 unique positions plus wild-type (WT), returning scores, assignments, and histograms for all. The “large” version of the workbook returns scores and assignments for the smaller of either all substitutions at 350 positions or 6000 substitutions. The first worksheet of each workbook contains detailed instructions on its use.
2. A Python package. The “rheoscale” python package generates scores, histograms, and assignments for large datasets. It can be integrated into pipelines for analyzing HTP data, supports integration with Jupyter notebooks, and is accompanied by a readme file that describes its implementation. This program has three dependencies: numpy^26^ and pandas^27^, and matplotlib^28^. The Python implementation of RheoScale 2.0 is available through PyPI and can be installed using Python’s package manager, pip (https://pypi.org/project/rheoscale/). Unlike the Excel workbooks, the Python script has no limit on the number of substitutions and positions that can be analyzed; its runtime and maximum dataset size will depend on available computational resources. Both Excel and Python implementations can be found at https://github.com/liskinsk/RheoScale-calculator.

### Definitions and hierarchy of position classes

1. In RheoScale 2.0, the first decision is to determine whether a position is <u>neutral</u> for the measured feature. The default definition is that ≥70% of these positions’ substitutions have WT-like outcomes, using the “neutral bin” described in the next section. The rationale for this default threshold was extensively discussed in Martin *et al*.^25^ As further described below, the default threshold can be changed by the user when deemed appropriate.
2. Next, the neutral bin is used to assess how many substitutions are “better” than WT. If ≥80% of a position’s substitutions have values “better” than the neutral range, this position is defined as <u>enhancing.</u>^6^ To date, we have only encountered a few enhancing positions and the biological implications of their existence is not yet clear. One possibility is that a trade-off between two protein features constrained the evolution of the natural protein. For example, a position classified as enhancing in a stability assay may be a toggle position for activity and, consequently, would be selected against during evolution. Alternatively, the WT protein could also be “caught” at an unusual evolutionary step, after a recent change created new enhancing possibilities that have not yet been explored. The remaining non-neutral positions are further classified according to the following logic:
3. If most of a position’s substitutions lack detectable signals, it is designated as a <u>toggle</u> position. The default threshold in the calculator, 64%, was derived from a database for *Escherichia coli* lactose repressor protein.^5; 29^ This suggested threshold was chosen to allow physiochemically “similar” amino acids to substitute for each other, as has been experimentally observed for positions that are intolerant to most substitutions. Again, the default threshold can be changed by the user when deemed appropriate. Our choice to assign toggle positions before rheostat positions meets the logical requirement of assigning the extreme outcomes (toggle and neutral positions) to provide the context needed for assigning positions with the intermediate outcomes (*e.g.,* rheostat positions). As such, in the hierarchy of position assignments, toggle assignments take precedence even if the remaining substitutions surpass the rheostat score threshold described below.
4. The set of substitutions for a <u>rheostat</u> position widely samples the range of possible values for the measured feature. This is quantified via the rheostat score, which is calculated by scoring occupancy of each histogram bin as empty (0) or filled (1), regardless of how many substitutions occupy the bin. Simplistically, the set of substitutions at rheostat positions sample at least half of the possible range;^23^ the use of weighted rheostat scores privilege substitutions that differ most from both the WT and inactive variants.^23^ Weighted rheostat scores (described in Hodges *et al.*^23^ and the instructions that accompany the RheoScale 2.0 calculators) are used in the assignment hierarchy with a default threshold of 0.5. In addition to the three original assignments defined using the thresholds described above (rheostat, toggle, and neutral), two intermediate behaviors have been observed in experimental datasets:
5. At a <u>moderate</u> rheostat position, its set of substitutions has non-neutral outcomes, but (i) sample less than half of the possible range and (ii) the average of the experimental values are closer to the WT than to inactive variants.^8^
6. <u>Adverse</u> positions, like moderate positions, have substitutions with non-neutral outcomes that sample less than half of the possible range. However, the average outcome for their sets of substitutions are closer to the inactive value than to WT.^6^ The existence of these six position classes indicates that, in addition to the continuum of outcomes available from single substitutions at rheostat positions, the positions in a protein can fall along a continuum of position classes. Efforts are underway to discriminate the biophysical bases of the different position types (*e.g.,* ^17; 18; 30^). In future studies, it will be particularly interesting to determine whether the biophysical characteristics of moderate positions falls on the continuum between neutral and rheostat positions and whether those of adverse positions are between rheostat and toggle positions.
7. Finally, some positions have shown a “<u>WT/Inactive split</u>” behavior. That is, around half of their substitutions are like WT and the other half lack detectable signal; very few show intermediate outcomes.^24^ This assignment may be a hallmark of altered protein stability.^31^

### Key analysis parameters

For RheoScale 2.0 analyses, the user must balance considerations arising from: the numbers of variants available for each position, the error of the experimental data, the width of the neutral bin, the total range of the histogram, the number of bins used to calculate the rheostat score, and the score thresholds used to make assignments. Although the calculator uses the input data to recommend default values as reasonable starting places for various parameters, these parameters should be systematically varied for each dataset to assess the robustness of position assignments. Edge cases – which have been rare in our prior studies – require user interpretation.

Detailed descriptions of the <u>effects of experimental error</u> on the calculations and position assignments are described in ^23; 25^. Additional discussions of the other parameters are in the next paragraphs.

1. To identify rheostat positions, experimental data should contain <u>multiple variants</u> for each amino acid position analyzed. Although 19 substitutions (plus WT) per position are ideal, we previously reasoned that 10 substitutions per position should be sufficient to estimate the locations of rheostat and toggle positions.^23^ We also identified rheostat positions with as few as 5 variants, reasoning that since the available set sampled more than half the histogram range, having more variants would not change the rheostat assignment.^18^ Unambiguous assignment of neutral positions requires many variants per position and assays of multiple protein features; it is hard to prove a negative.^25^ To further assess the relationship between the number of available variants and the robustness of RheoScale 2.0’s assignments, we here used random sampling of the PLpro HTP activity dataset.^12^ The PLpro library contains variants for 315 positions. Of these, 96 positions have assay results for all 20 variants (19 substitutions plus WT), 190 positions have results for 15-19 substitutions, 29 have results for 10-14 substitutions, and 2 positions have results for 7 substitutions. For this exercise, we selected the positions with all 20 substitutions and used random sampling to generate 15 subsets with 5-19 substitutions per position. For each subset, random sampling was repeated ten times, and two RheoScale analyses were carried out: (i) using ten histogram bins (Figure 2, filled squares) and (ii) using the default bin number recommended by Rheoscale 2.0 (Figure 2, black, open circles). The assignment accuracies were then calculated relative to assignments using all 20 variants for each position. For each set of ten random replicate subsets, we then calculated average accuracies and standard deviations (Figure 2; https://github.com/liskinsk/RheoScale-calculator/tree/main/Plpro%20HTP%20examples). As expected, assignment accuracy increased with the number of available variants. Nevertheless, when 10 histogram bins were used, assignments approached 90% accuracy with only 13 variants per position and >70% of the changed assignments were to an adjacent category (*e.g.,* a rheostat assignment changed to either an adverse or a moderate assignment) for all sample sizes. Using the RheoScale 2.0 default-recommended bin numbers, the missed assignments included more non-adjacent categories; however, there were fewer incorrect assignments overall. Even 7 variants per position showed >80% accuracy when the RheoScale 2.0’s recommended bin number was used (Figure 2).
2. The <u>width of the neutral bin</u>, which is distinct from the histogram bins, is the key parameter for assessing whether variants are equivalent to WT and, in turn, whether a position is neutral.^25; 32^ The neutral bin is centered on the value measured for the WT protein; the default width of the neutral bin width is +/-2 standard deviations of the WT error (or four times the error override for the full dataset). If no error is designated, the neutral bin width is set to twice the width of the histogram bin that is used to calculate rheostat and toggle scores. The calculator also provides an opportunity for the user to manually define the neutral bin size. Variants with values that fall in the neutral bin are designated as “WT-like”.
3. The most important parameter for assigning non-neutral positions is the <u>histogram range</u>. In RheoScale 2.0, the range defaults to the minimum and maximum of the experimental data. If experimental data do not reflect the actual range, *e.g.* the measured outcomes only sample a portion of the assay’s accessible range, the user can enter override values. Various examples that require user-determined values, including strategies for assigning “dead” values that define the bottom of the range, are in the documentation that accompanies the RheoScale 2.0 calculators.
4. The recommended <u>number of histogram bins</u> is calculated from the average error for the dataset and the number of variants available for each position in the dataset.^23^ Since rheostat positions are defined by having substitutions with intermediate loss of signal, the resolution of the experimental data is critical. Ideally, at least three histogram bins will contain intermediate values. The user can also choose a bespoke bin number; empirically, 10 histogram bins work well for many HTP datasets. Indeed, iterating RheoScale analyses with different bin numbers has shown that most position assignments are not very sensitive to this parameter (*e.g.,* ^33^): For a given histogram range, decreasing the bin number increases the bin width. Although this can be appropriate for accommodating larger experimental errors, it is also more likely that a set of substitutions will sample a large fraction of the available bins. These two features usually offset each other, resulting in the same position assignment. This was seen for assignments using activity values for the full set of PLpro positions (with 7-20 variants) and with bin numbers ranging from 7 to 14 (*e.g.,* Figure 3). The exception is sparse datasets (*e.g.,* Figure 2; 5-7 variants per position), where using the smaller, RheoScale-recommended number of bins has better accuracy than a larger number that forces several bins to be empty (*i.e.,* five variants can never fully sample 10 bins.). Of the PLpro positions that changed assignment classes in this exercise (51 of 315, which is 16%), the majority changed to an adjacent category (*e.g.*, rheostat to moderate). This is expected when a continuous property is subjected to thresholds^34^, which are here imposed by bin boundaries and assignment score thresholds. At the largest bin number used in this exercise (14), most of the reassignments were rheostat positions to moderate or adverse positions. We considered whether the reassigned positions were dominated by positions with <14 variants, but the fraction of changed assignments in this subset (5 of 20, or 25%) was only marginally higher than the overall fraction of changes (16%). This suggests that most reassignments occur when substitution values fall near the boundaries of a bin (which changes with bin number). For the PLpro activity results presented below, we used 10 bins to balance the competing interests of (i) accommodating experimental error (requiring fewer histogram bins) and (ii) avoiding over-predicting the number of rheostat positions (via more histogram bins). For other datasets, another strategy would be to vary the bin number and choose the consensus assignment for each position.
5. For sparse or low resolution datasets, <u>assignment score thresholds</u> should also be considered. Stricter thresholds (such as requiring 75% of variants to be inactive when assigning toggle positions) provide more confidence in position assignments, especially for sparse experimental datasets (*e.g.*,^8; 18^). As with bin numbers, assignments should be iteratively evaluated for their robustness to different thresholds; position assignments that are not robust should be manually inspected.

**Figure 2.**
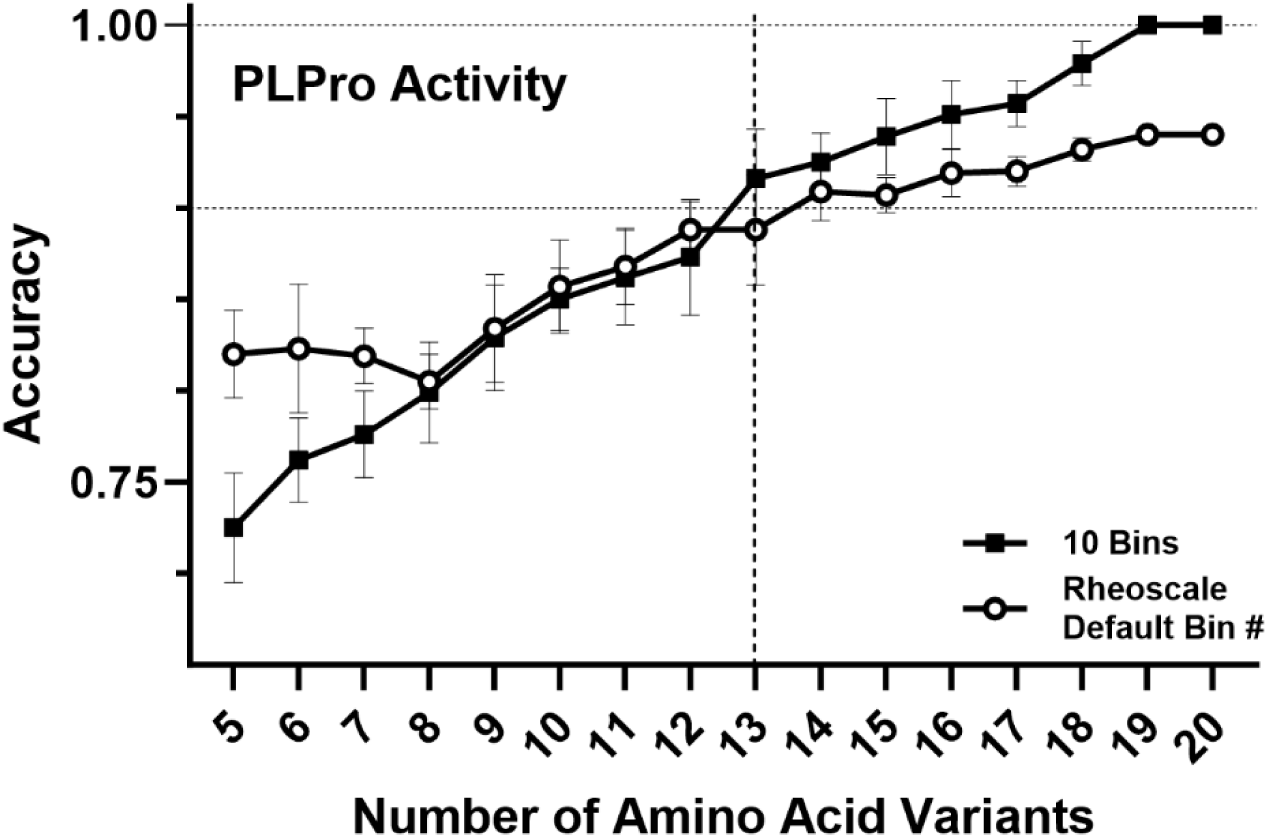
RheoScale 2.0 assignments are robust to changing variant numbers. To assess how the number of available variants alters position assignments, we used positions in the HTP dataset of PLpro activity for which all twenty variants are available. The variant set for each position was randomly subsampled to generate 10 replicates for each of 5-20 variants. RheoScale analyses were carried out for each subsample using either 10 bins (squares) or the default number of bins recommended by RheoScale (open circles); note that the recommended default bin number is based on the number of available variants, as described in ^23^, and thus differs with variant number. For each variant number, assignment accuracy was compared to the assignments using all 20 variants and 10 bins as “truth”, and the average and standard deviation for each set of subsamples is plotted. All class assignments used default score thresholds. Dashed lines are to aid visual inspection of the data.

**Figure 3.**
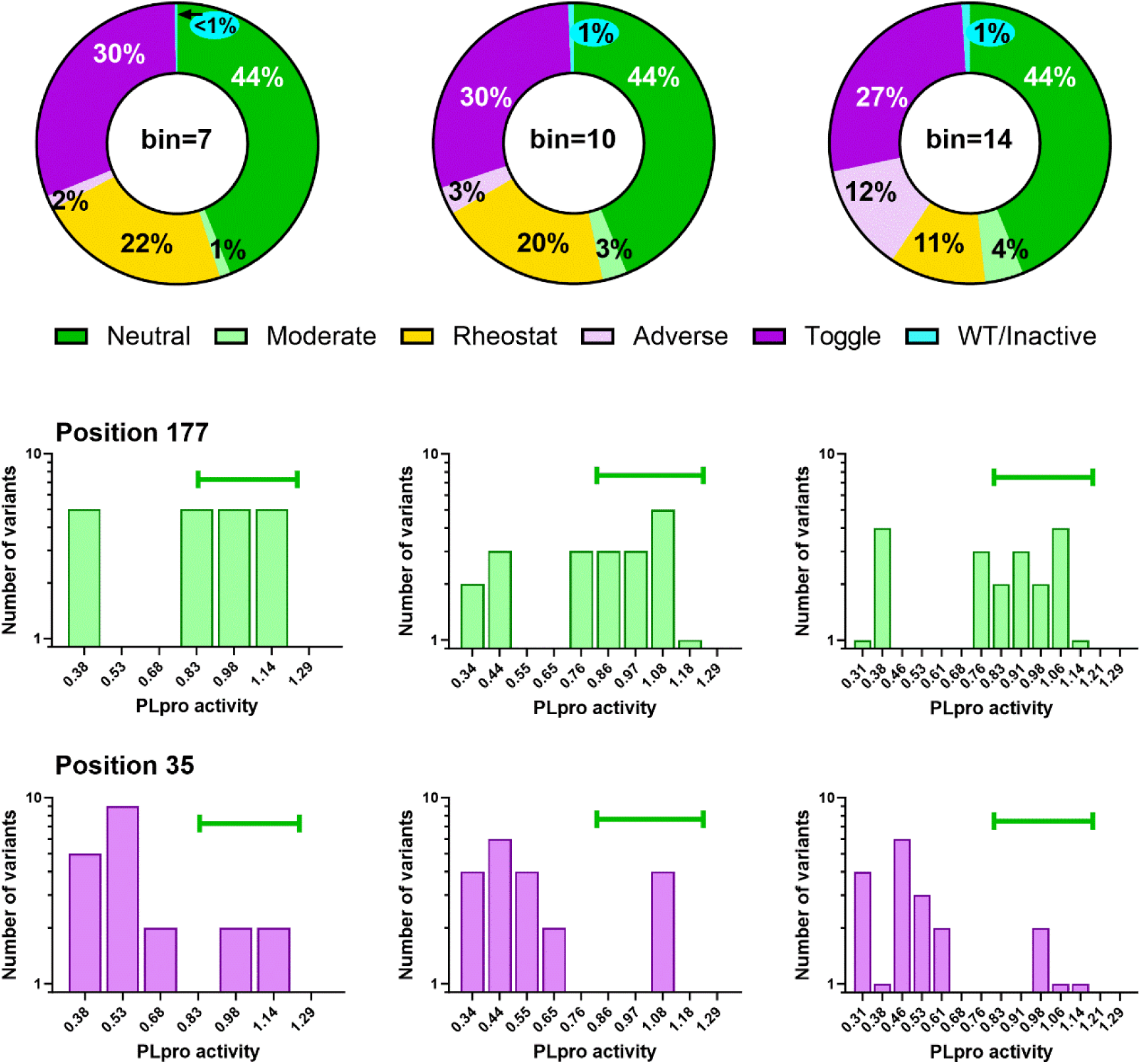
Effects of changing bin numbers on position assignments. RheoScale analyses were carried out for the PLpro activity data^12^, with bin numbers ranging from 7 to 14. Position classes are colored as follows: neutral, green; moderate, light green; rheostat, yellow; adverse, light purple; toggle, purple; and WT/inactive split, cyan. Histogram bins are labeled with their upper limit, as in Figure 1. The most frequent assignment changes were when rheostat positions changed to an adjacent category (either adverse or moderate). The histogram plots show examples for this behavior using 7, 10, and 14 bins: position 177 changes from moderate to rheostat and back to moderate; position 35 changes from rheostat to adverse and back to rheostat. Note that the fraction of neutral scores does not change because these assignments are made using a separate “neutral bin” (green horizontal bar on the histograms).

## Results and Discussion

To demonstrate the utility of RheoScale 2.0, we used three deep mutational scanning datasets for a site-saturating mutagenesis library of SARS-CoV-2 PLpro.^12; 14^ (i) The activity assay measured *in vivo* proteolysis of a fluorescent reporter protein after PLpro expression was induced in mammalian cells.^12^ (ii) Changes in *in vivo* abundance were measured using a construct that fused PLpro directly to a fluorescent reporter protein.^12^ (iii) Finally, another implementation of the activity assay included enzyme inhibitors; the dataset used here was generated by the original authors by combining HTP assays using two different inhibitors.^14^ Analyses of these datasets reveal diversity of position assignments, as observed in many other proteins.^7^ All RheoScale 2.0 assignments for PLpro, as well as a table compiling assignments from the three assays, are available at https://github.com/liskinsk/RheoScale-calculator.

In the PLpro protease activity data (Figure 4), ∼20% of positions were rheostat positions. Similar to other proteins,^7^ rheostat positions are dispersed across the structure, which underscores the fact that a position’s substitution sensitivity cannot be inferred from its location. As anticipated, RheoScale 2.0 analyses further parsed PLpro positions with similar average scores (*e.g.*, Figure 5) into different position classes. This is important because substitutions at rheostat positions appear to arise from special biophysical characteristics (*e.g.,* ^18^).

**Figure 4.**
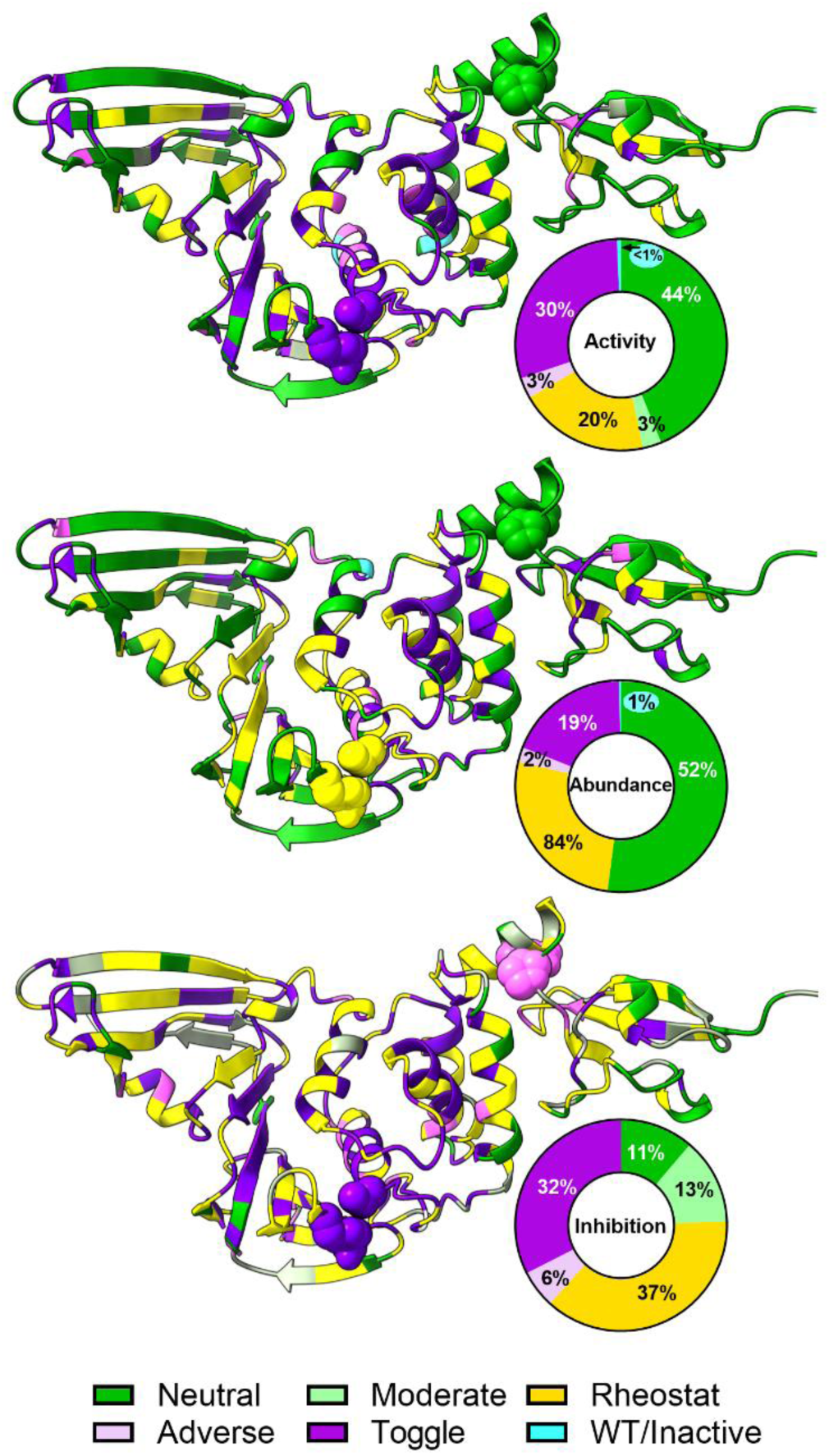
PLpro positions exhibit assay-dependent assignments. Ribbon representations of SARS-CoV-2 PLpro (PDB: 8VEC^12^) are color-coded according to their RheoScale 2.0 position assignments for (top) protease activity, (middle) *in vivo* abundance, and (bottom) activity in the presence of inhibitors. Donut charts summarize the fraction of PLpro positions assigned to each position class. Position classes are colored as follows: neutral, green; moderate, light green; rheostat, yellow; adverse, light purple; toggle, purple; and WT/inactive split, cyan. The catalytic triad residues C111, H272, and D286 are shown as purple spheres on the structure showing activity and inhibition data and yellow on the abundance data; these positions are classified as toggle in the activity and drug escape but rheostat in abundance. Positions 59 and 67 are also highlighted as spheres (pink on the structure showing inhibition assignments); these positions are near the domain interface and are neutral in the activity and abundance datasets but become adverse in the inhibitor dataset. Structure images were generated using UCSF ChimeraX.^40^

**Figure 5.**
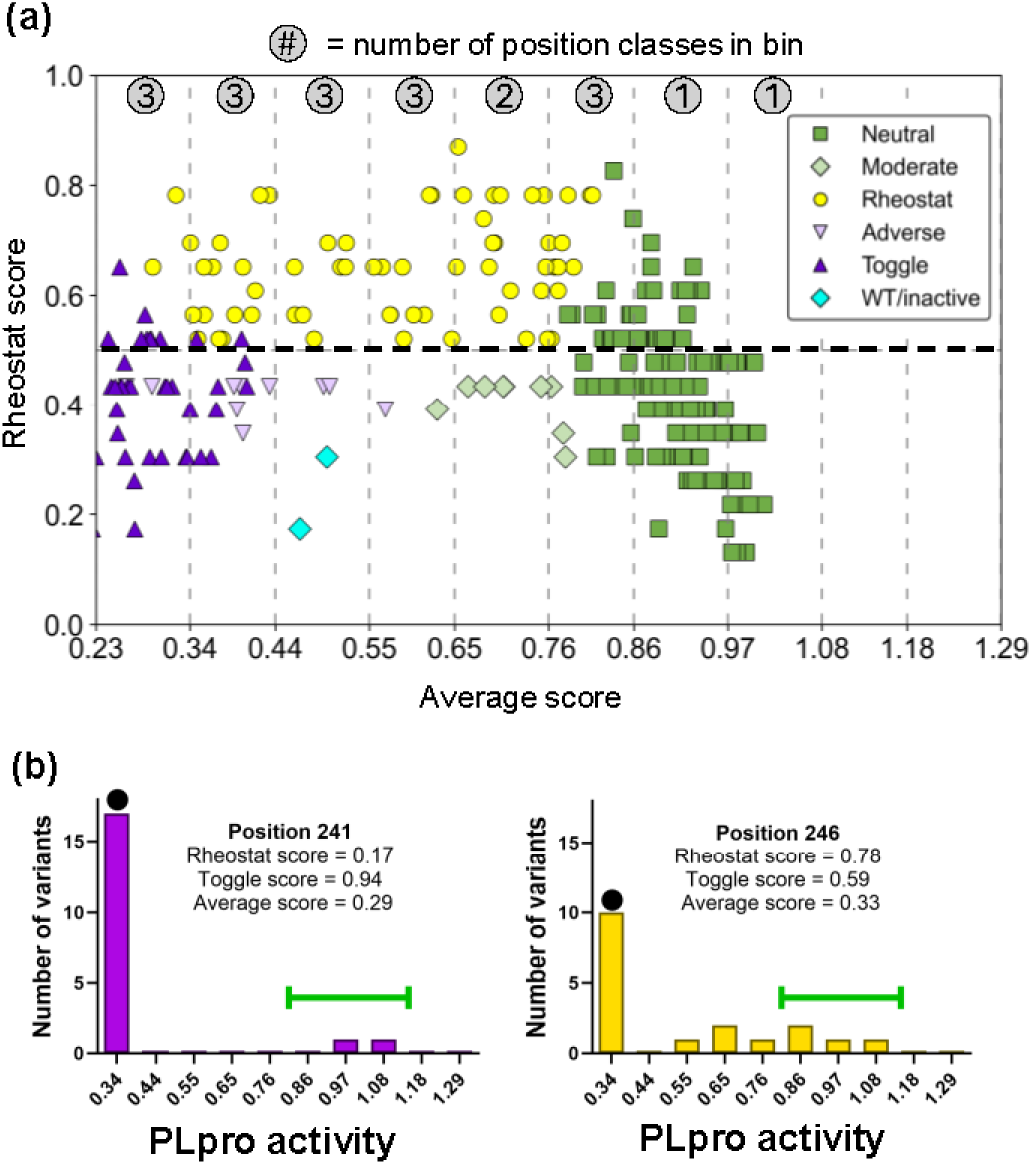
RheoScale 2.0 assignments using SARS-CoV-2 PLpro activity data^12^. (a) Rheostat scores versus average scores for each PLpro position. Each point on the XY plot represents a unique PLpro position; symbols correspond to position assignments. The dashed horizontal line indicates the threshold of 0.5 that is used to assign rheostat positions (after neutral and toggle assignments are made). The dashed vertical lines divide the average values into the same bins as in Figure 1, and the numbers in gray circles denote how many position classes are represented for each bin. An example of two positions with similar averages but different rheostat scores are shown in (c). Position 241 (left) demonstrates toggle behavior; position 246 (right) displays rheostat behavior. Other features of these plots are as described in Figure 1. In (b) and (c), the values on the x-axes correspond to the upper bin limits.

Comparing results for activity and abundance assays illustrates that each position on a protein’s scaffold can play specific roles. For example, PLpro positions that are classified as toggle positions for both activity and abundance may primarily contribute to protein stability. Other positions show no effect on abundance but do show effects on activity, suggesting that these positions contribute primarily to function. Still other positions affect both features, and as observed for Mpro^6^, the altered activity may arise from simultaneously changing both function and abundance. Additional “layers” of position roles might be expected if PLpro were to be assayed for different activities such as de-ubiquitinylation and de-ISGylation of host proteins; these functions use the same active site but different substrate binding sites.^35^ We have observed similarly complex relationships between position roles and distinct functional features in several other proteins.^7; 8; 31–33^

The comparison of activity and abundance data also highlights the importance of understanding assays detection limits. In general, one would expect that a change in activity would be constrained by any change in abundance. That is, substitutions with undetectable abundance should not have activity, and thus positions that are toggle for abundance should not be rheostatic for activity. However, some PLpro positions that are classified as toggle for abundance are rheostat positions for activity. A plausible explanation for this apparent paradox is that the activity assay is more sensitive than the abundance assay. This difference precluded calculations to dissect functional information from the activity score (*e.g.,* ^6; 33; 36^), which is a composite of both function and abundance.

When results for the inhibitor assay are compared to the activity assay, many PLpro positions show greater sensitivity to substitution and more are classified as rheostat positions (37% vs 20%; Figure 4). This can be explained by the inhibitor lowering the effective *in vivo* concentration of active PLpro for the majority of substitutions. This observation also suggests that the first activity assay – with data collected at a single time point 24 hours after PLpro induction – was performed under “V_max_” conditions that obscured the effects of many substitutions. Similar findings were observed from comparisons of two Mpro HTP datasets assayed at different induction times.^6; 10; 37^

In addition, one might expect some PLpro substitutions to be more or less sensitive to inhibitors than others. Indeed, Call and colleagues identified five positions with substitutions resistant to inhibitor^12^; four of these positions (PLpro 164, 167, 208, and 269) are classified as rheostat positions in the inhibitor assay, while the fifth (268) is classified as a moderate position. These five positions were also the locations of resistance mutations for more potent inhibitors ^14^, as were positions 266 and 267, which are also rheostat positions in the weak inhibitor assay; a third position for an escape mutant (301) was a toggle position. Thus, 75% of the positions with escape mutants were rheostat positions, which is essentially double the probability of sampling a rheostat position by random chance (37%; see Figure 3 legend). This suggests that amino acid changes at rheostat positions can play important roles in viral evolution and drug escape, and identification of rheostat positions could be beneficial for focusing further studies.

RheoScale 2.0 analyses also identify two positions with higher-than average sensitivity to inhibitors (PLpro positions 59 and 67). Intriguingly, these two positions are near each other on the structure and near a domain interface (Figure 4, pink spheres in inhibition data). Biochemical and biophysical studies of these and other rheostat positions should be fruitful for understanding the sequence-structure-function relationship of this PLpro construct at a molecular level.

Finally, we used the PLpro analyses to consider how different types of experimental data (*e.g.* Figure 4) correlate with predictions of position assignments. The funtrp algorithm predicts whether positions exhibit neutral, toggle, or rheostat behavior.^16^ This algorithm was trained on HTP datasets for five proteins, each of which measured substitution effects on a feature specific to that protein (or, in cell-based assays, a combination of features such as function and abundance). Additional input features included (i) a variety of biophysical characteristics about each amino acid and its predicted location in the protein structure, (ii) the number of possible single nucleotide polymorphisms that lead to a missense mutation, and (iii) ConSurf^38; 39^ scores, which are derived from phylogenetic conservation analyses of multiple sequence alignments. ConSurf scores contributed the most to funtrp predictions.^16^ Because of this, we chose to focus our comparisons on the PLpro activity and abundance measurements, reasoning that, since inhibitors were not part of PLpro’s natural evolution, they would not leave a “signature” in the multiple sequence alignment of PLpro homologs.

We also considered that, in training funtrp, the thresholds used for classifying neutral and toggle positions differed from the default RheoScale thresholds: Neutral positions were designated when “all but one” variants were wild-type.^16^ Toggle positions were designated when “all but two” variants were detrimental.^16^ Since the number of variants in the funtrp training dataset ranged from 6 to 20 per position, converting these to fraction-based thresholds comparable to those of RheoScale 2.0 spans a range: The neutral threshold could range from 0.80 to 0.95 and the toggle threshold could range from 0.67 to 0.9. Since the lower ends of those ranges are similar to RheoScale 2.0 default values, we repeated the comparisons using the stricter thresholds of 0.9 for both neutral and toggle assignments.

Our first question was how funtrp handles positions experimentally assigned to the moderate and adverse classes, since these were not described when funtrp was developed. Using RheoScale’s default score thresholds, the experimental abundance data had seven adverse positions; funtrp predictions for these positions were split among rheostat and toggle classes, both of which are reasonable assignments. In the activity data, five of the nine moderate and six of the ten adverse positions were assigned to the rheostat category, two moderates were predicted to be neutral, and three adverse were predicted to be toggle. Again, these are reasonable, “correct” predictions. However, two positions that were moderate for activity were predicted to be “toggle” and one adverse position for activity was predicted to be “neutral”.

Our second question was to examine the effects of stricter thresholds. Results showed that the number of positions that were assigned as neutral or toggle decreased in both datasets, as expected (Figure 6). Many of the default-neutral and -toggle positions were reassigned to the moderate and adverse classes, respectively, because variants that fell outside of the neutral/toggle bins were located in nearby bins. This is consistent with the emerging picture that individual positions fall along a continuum of substitution sensitivities. Other positions changed from neutral (or toggle) to rheostat assignments. This arose when the remaining variants widely sampled the rest of the histogram bins; that is, positions with five non-neutral (or non-toggle) variants could occupy enough bins to reach a rheostat score ≥0.5.

**Figure 6.**
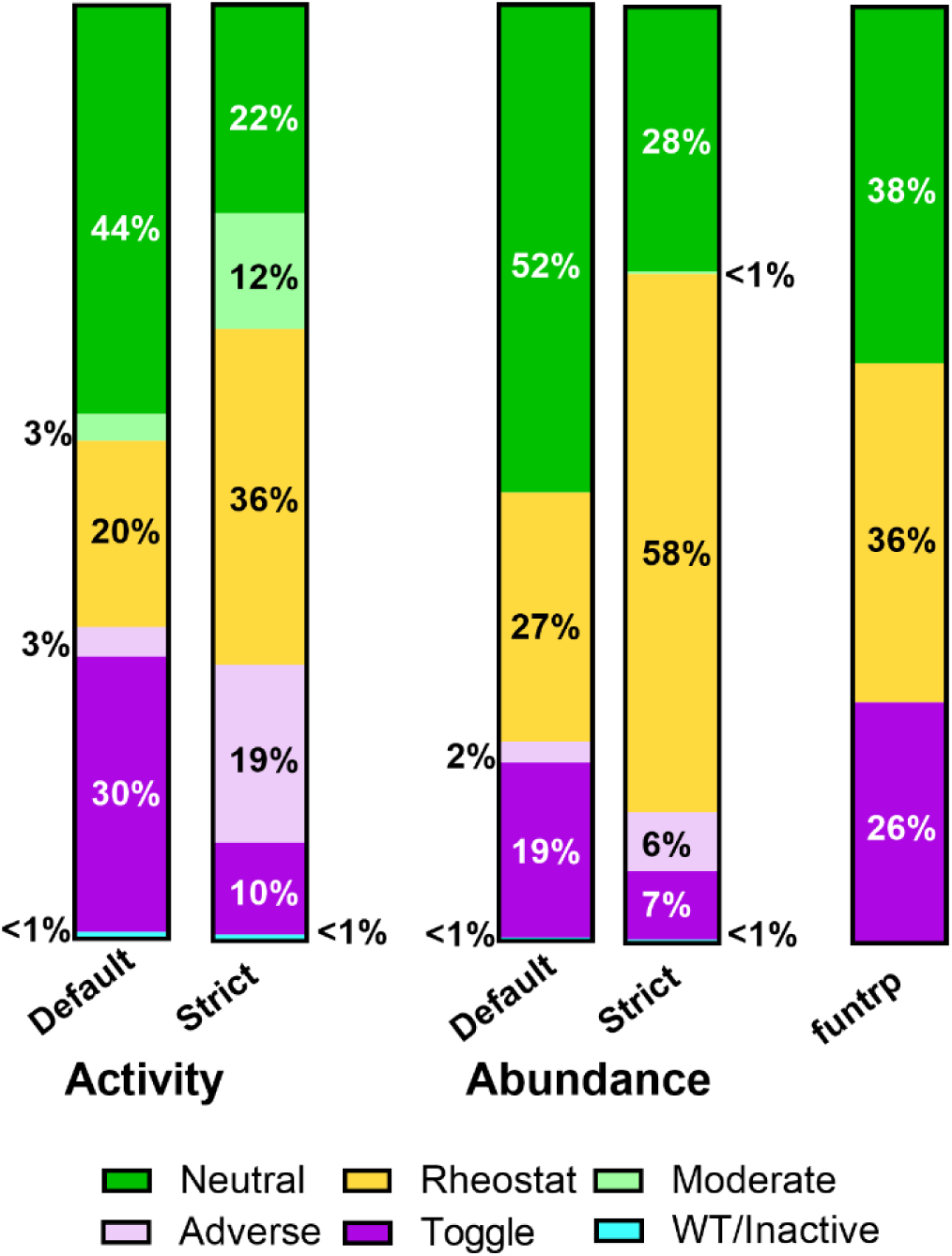
Comparison of PLpro experimental and predicted position classes. The percent of positions in the PLpro activity and abundance datasets that were experimentally assigned to each class were determined using (i) RheoScale 2.0 default or (ii) strict toggle/neutral thresholds. For the activity data, 30 positions changed from neutral to moderate, 50 changed from toggle to adverse, 38 changed from neutral to rheostat, and 12 changed from toggle to rheostat; 101 neutral and toggle position assignments stayed the same. For the abundance data, 1 position changed from neutral to moderate, 13 changed from toggle to adverse, 74 changed from neutral to rheostat, and 23 changed from toggle to rheostat; 112 neutral and toggle assignments stayed the same. The greater number of rheostat re-assignments in the abundance data is because RheoScale analyses used fewer histogram bins (seven) than the activity data (ten); this was due to the lower experimental resolution of the abundance data. The right-most bar shows the predicted distribution of positions among the three funtrp classes (neutral, rheostat, and toggle).

Our third question was related to the overall success of funtrp for the two datasets. For these comparisons, the experimentally-moderate and -adverse assignments were re-classified as rheostat positions. Using the RheoScale default thresholds, funtrp’s overall prediction success was greater than 50% for both experimental datasets, which exceeds random chance (33% for a three-category assignment). We also considered that the mismatch could arise from either a false prediction or an experimental misclassification. However, there was no correlation between prediction accuracy and the number of variants available to make the experimental assignment, reducing the likelihood that a false prediction arises from a false experimental assignment. Instead, prediction accuracy differed among position classes, with experimentally-neutral and -toggle positions better predicted than rheostat positions (Table 1).

**Table 1.**
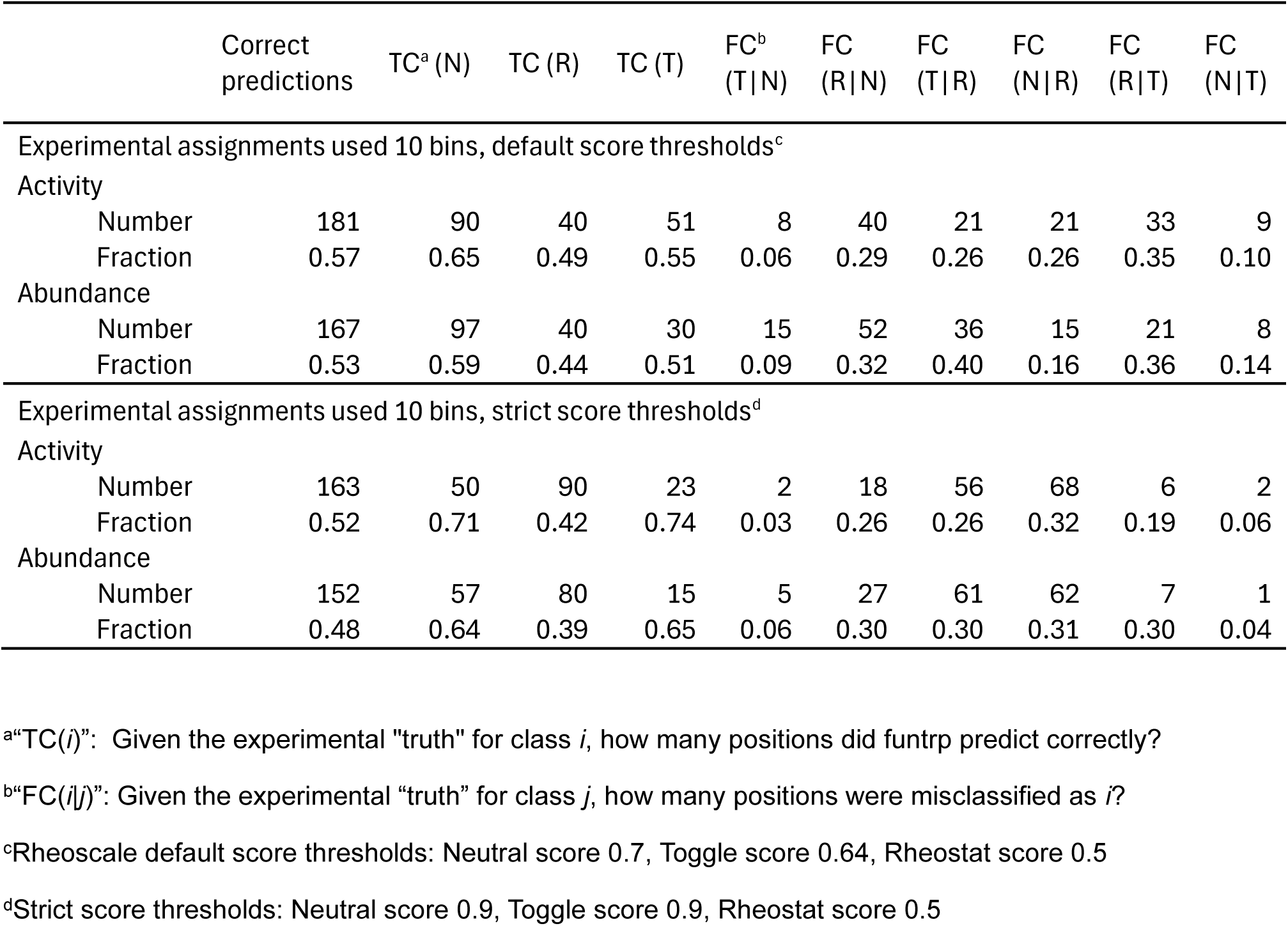
Statistical analyses of funtrp predictions for PLpro position classes. ^a^“TC(*i*)”: Given the experimental “truth” for class *i*, how many positions did funtrp predict correctly? ^b^“FC(*i*|*j*)”: Given the experimental “truth” for class *j*, how many positions were misclassified as *i*? ^c^Rheoscale default score thresholds: Neutral score 0.7, Toggle score 0.64, Rheostat score 0.5 ^d^Strict score thresholds: Neutral score 0.9, Toggle score 0.9, Rheostat score 0.5

|  | Correct<br>predictions | TC<sup>a</sup> (N) | TC (R) | TC (T) | FC<sup>b</sup><br>(T\|N) | FC<br>(R\|N) | FC<br>(T\|R) | FC<br>(N\|R) | FC<br>(R\|T) | FC<br>(N\|T) |
| --- | --- | --- | --- | --- | --- | --- | --- | --- | --- | --- |
| Experimental assignments used 10 bins, default score thresholds<sup>c</sup> |  |  |  |  |  |  |  |  |  |  |
| Activity |  |  |  |  |  |  |  |  |  |  |
| Number | 181 | 90 | 40 | 51 | 8 | 40 | 21 | 21 | 33 | 9 |
| Fraction | 0.57 | 0.65 | 0.49 | 0.55 | 0.06 | 0.29 | 0.26 | 0.26 | 0.35 | 0.10 |
| Abundance |  |  |  |  |  |  |  |  |  |  |
| Number | 167 | 97 | 40 | 30 | 15 | 52 | 36 | 15 | 21 | 8 |
| Fraction | 0.53 | 0.59 | 0.44 | 0.51 | 0.09 | 0.32 | 0.40 | 0.16 | 0.36 | 0.14 |
| Experimental assignments used 10 bins, strict score thresholds<sup>d</sup> |  |  |  |  |  |  |  |  |  |  |
| Activity |  |  |  |  |  |  |  |  |  |  |
| Number | 163 | 50 | 90 | 23 | 2 | 18 | 56 | 68 | 6 | 2 |
| Fraction | 0.52 | 0.71 | 0.42 | 0.74 | 0.03 | 0.26 | 0.26 | 0.32 | 0.19 | 0.06 |
| Abundance |  |  |  |  |  |  |  |  |  |  |
| Number | 152 | 57 | 80 | 15 | 5 | 27 | 61 | 62 | 7 | 1 |
| Fraction | 0.48 | 0.64 | 0.39 | 0.65 | 0.06 | 0.30 | 0.30 | 0.31 | 0.30 | 0.04 |

When we used stricter score thresholds for assigning experimentally-neutral and -toggle positions, funtrp prediction accuracy increased for these two classes (Table 1). However, the accuracy for predicted rheostat positions decreased (Table 1), as did the overall accuracy. This could be because the toggle and neutral positions with the most extreme substitution sensitivities also have the most extreme conserved and non-conserved ConSurf scores, respectively.^8; 25^ Given the importance of ConSurf scores to funtrp predictions, the funtrp toggle positions may be among the most conserved PLpro positions; likewise, the funtrp neutral positions may be among the least conserved.

Thus, positions with intermediate conservation remain among the hardest to predict for both position classes and substitution outcomes. We hypothesize that the discrepancies might be explained by our previous findings in a range of proteins, where outcomes at rheostat positions were not well-explained by the common structural properties^7^ that are used as other input in funtrp. One possible route for improving predictions is the addition of more protein features; for example, we found that information about coupled dynamic motions were able to discriminate neutral and rheostat positions in the lactose repressor protein.^18^ Alternatively, the multiple sequence alignment created by ConSurf could be improved for PLpro via manual curation (which is not currently an option in funtrp); in our hands, we have found that the over-representation of SARS-CoV2 sequences makes it difficult for automated searches to identify the broadest possible range of homologs.

Finally, we note that all activity classes were better predicted than abundance classes (Table 1). This is consistent with the fact that both the activity measurements and the evolutionary information used in funtrp/ConSurf are sensitive to both changes in catalytic activity and abundance. As such, funtrp – which is trained on fitness data that likely contains information from variant effects on both function and abundance – might be more suitable for predicting the composite sensitivity of each position rather than position roles in specific biochemical features, such as binding affinities, allosteric response, or stability.

## Conclusion

By capturing both the extreme and intermediate substitution outcomes, RheoScale 2.0 reveals position-specific behaviors that are obscured by average values (Figure 5). Understanding these intermediate outcomes, and the rheostat positions that host such changes, can help prioritize research for mechanistic studies, protein engineering, and drug development. For example, the enrichment of inhibitor-resistance substitutions at rheostat positions in PLpro suggests that substitutions at rheostat positions provide accessible routes for viral evolution and drug escape.

The current results demonstrate that RheoScale 2.0 assignments (i) are robust to variations in subjective analysis parameters and (ii) that position assignments are >90% accurate with as few as 13 variants per position, which can ameliorate the costs and time required to generate all 19 variants for every position. The experimental error, the dynamic range of the assay, and data resolution remain important considerations for revealing the evolutionarily-important intermediate outcomes at rheostat, moderate, and adverse positions.

The PLpro example that we used to carry out these statistical assessments of RheoScale 2.0 analyses also illustrates an important characteristic of the protein sequence-structure-function relationship: A given amino acid position can have more than one role in the protein architecture, and the position can have a different substitution sensitivities for its different roles (*i.e.,* catalytic activity, abundance/stability, or inhibitor response). Thus, it is imperative that researchers recognize the feature(s) detected by each assay and understand when or if some substitution effects are concealed by assay design.

In summary, RheoScale 2.0 provides an objective framework for extracting position specific information from high-throughput datasets in both Excel and Python formats. Applying the framework across multiple assays and proteins will deepen our understanding of sequence-structure-function relationships, thereby improving our abilities to interpret pathological variants and identify positions well-suited for protein engineering and drug development.

## Credit authorship contribution statement

**David H. Liu**: Conceptualization, Writing – review & editing, Writing – original draft, Methodology, Formal analysis, Software, Investigation, Validation, Visualization. **Shwetha Sreenivasan:** Writing – review & editing, Writing – original draft, Methodology, Formal analysis, Conceptualization, Software, Investigation, Validation, Visualization. **Carter J. Gray:** Methodology, Writing – review & editing, Formal analysis, Conceptualization, Software, Investigation, Validation, Visualization. **Hannah C. Cleveland**: Methodology, Writing – review & editing, Software, Investigation, Validation. **Liskin Swint-Kruse:** Writing – review & editing, Writing – original draft, Formal analysis, Funding Acquisition, Conceptualization, Project administration, Supervision, Methodology, Data curation, Investigation, Validation, Visualization, Resources

## Acknowledgements

We thank Sara Volz (Northwestern University) for beta testing the Python version of RheoScale 2.0, Brett Gilio (KUMC) for assistance debugging the code, and Harshitha Vasan (KUMC) for preliminary analyses of PLpro. We thank Drs. Melissa Call and Xinyu Wu (Walter and Eliza Hall Institute of Medical Research) for discussions about the PLpro datasets. This work was supported by the National Institutes of General Medical Sciences [grant number R01GM147635 to LSK, P20GM103418 to CJG and HCC, and T34GM136453 to CJG].

## Declaration of generative AI and AI-assisted technologies in the manuscript preparation process

During the preparation of this work, we used Microsoft Copilot and ChatGPT to assist with the syntax of Excel formulas, reverse engineer the previously-published R version of RheoScale, and troubleshoot Python code. After using these tools, we reviewed and edited the output and take full responsibility for the content of the published article.

